# Comparative Study of BBB-Targeting AAV Capsids on Central Nervous System Delivery Efficiency

**DOI:** 10.64898/2025.12.23.696327

**Authors:** Junhao Zhao, Xiangyu Ge, Minghui Song, WeiMin Liu, Xia Zhang, Li Zuo, Lei Jin

**Affiliations:** Laboratory of Molecular Biology, Department of Biochemistry, School of Basic Medical Sciences, Anhui Medical University, Hefei, Anhui, China; Innovation and Entrepreneurship Laboratory for College Students, Anhui Medical University, Hefei, Anhui, China; Lingang Laboratory, Shanghai 200031, China; School of Life Science and Technology, ShanghaiTech University, Shanghai 201210, China; Experimental animal center, Anhui Medical University, Hefei, Anhui, China; Key Laboratory of Anti-inflammatory and lmmune Medicine (Anhui Medical University), Ministry of Education, Hefei, 230032, China

**Keywords:** PHP.eB, CNSRCV300, BI-hTFR1, Constitutive Promoter, Neuron-specific Promoter

## Abstract

The efficacy of adeno-associated virus (AAV)-mediated systemic gene therapy for central nervous system (CNS) diseases is often limited by the blood-brain barrier (BBB). This study systematically evaluated the tissue distribution of three BBB-crossing AAV capsid variants (PHP.eB, CNSRCV300, and BI-hTFR1) following intravenous injection in mice, using either a constitutive promoter (CAG) or a neuron-specific promoter (hSyn) to drive EGFP reporter expression. Compared with AAV9, both PHP.eB and CNSRCV300 demonstrated significantly enhanced BBB penetration and brain transduction efficiency. While the use of the hSyn promoter led to reduced transgene expression in the brain compared with the CAG promoter, and substantially decreased visible reporter expression in peripheral organs, viral deposition in the liver could still be detected via immunohistochemistry. Overall, CNSRCV300 exhibited the most favorable balance between brain-targeting efficiency and biosafety, highlighting its potential as a promising delivery vector. In summary, both the capsid and promoter jointly influence AAV-mediated expression *in vivo*, and although cell type-specific promoters can reduce off-target expression, residual viral deposition in non-target tissues remains a potential safety concern.

## Introduction

Adeno-associated virus (AAV) has been utilized as a gene delivery vector in the field of gene therapy for decades, owing to its inherent non-pathogenicity and long-term stability of gene expression. AAV is typically administered via intracerebral injection for *in vivo* gene delivery; however, this approach carries significant invasive risks. Moreover, due to the limited diffusion of viral particles within brain tissue, this method is generally effective only for treating focal lesions and often fails to achieve comprehensive coverage of whole-brain or extensive central nervous system pathologies[1, 2]. For patients with multifocal neurological disorders, repeated intracerebral injections at multiple sites are required, which increases both treatment burden and risks[3, 4]. While intravenous administration can ensure uniform gene expression in tissues, the presence of the blood-brain barrier severely restricts the entry of AAV vectors from the peripheral circulation into the central nervous system[5]. Since the initial report on the AAV-PHP.eB vector, a series of AAV capsid variants with enhanced blood-brain barrier penetration capabilities have been developed[6]. These engineered capsid variants typically achieve efficient transport from the peripheral blood to the brain parenchyma through receptor-mediated transcytosis via specific receptors on vascular endothelial cells[7-10].

This study describes three adeno-associated virus (AAV) capsid variants capable of crossing the blood-brain barrier (BBB)—PHP.eB, BI-hTFR1, and CNSRCV300. These variants differ in their amino acid sequences within the VP-VIII region near residue 588 (Fig. 1a).

**Fig 1.**
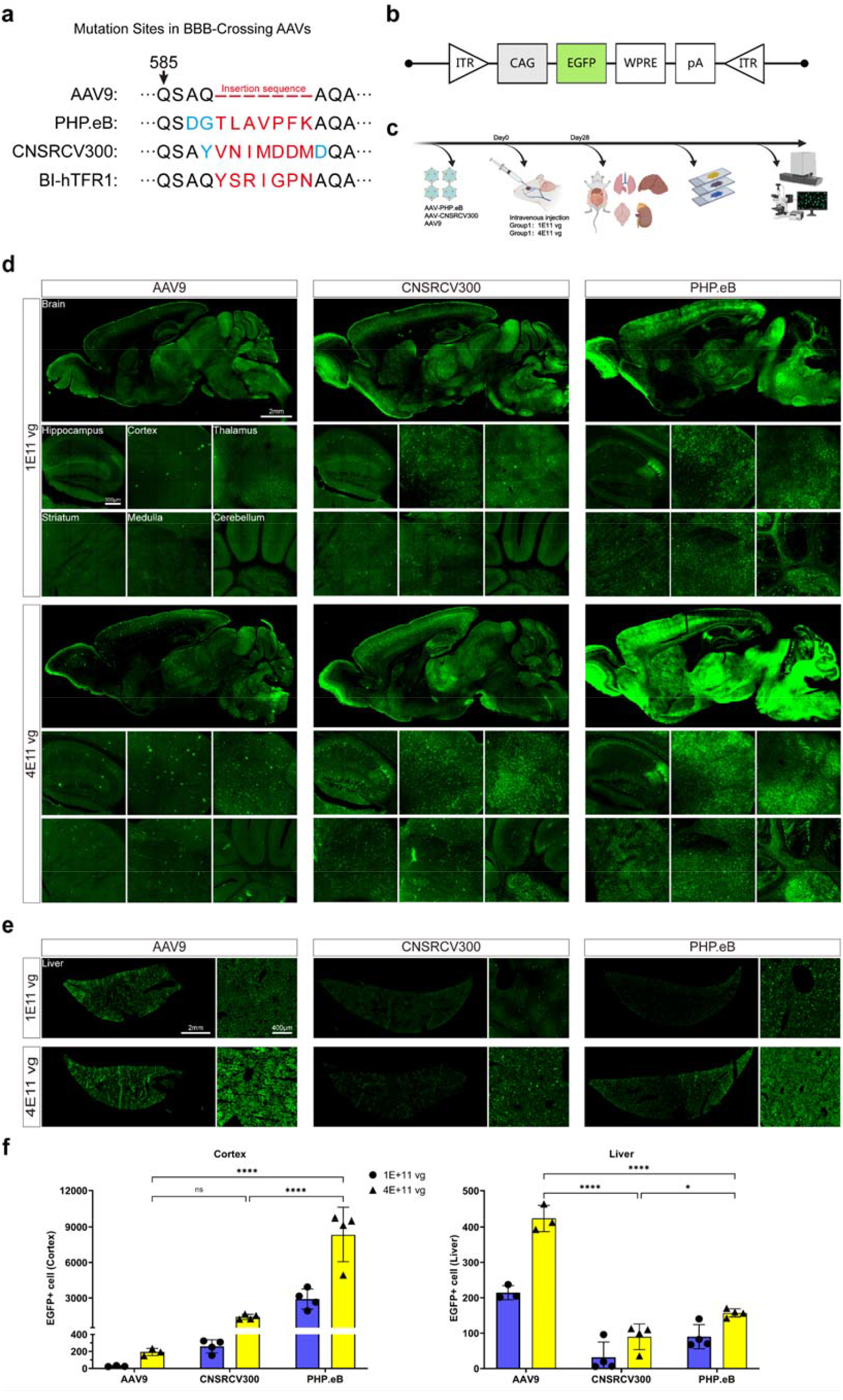
Transgene expression mediated by AAV9-derived capsids carrying the CAG promoter in brain and liver tissues. (a) Schematic representation of amino acid residue differences among the four capsid variants (AAV9, PHP.eB, CNSRCV300, BI hTFR1) that were used in the paper. (b) Structure of the transgene plasmid containing the constitutive CAG promoter, EGFP, WPRE and polyA elements. (c) Experimental timeline: PHP.eB and CNSRCV300 viruses were intravenously injected into adult C57BL/6J mice at two doses (1.0×10^11^ vg and 4.0×10^11^ vg; n = 3–4 per group). Tissues were collected and sectioned 4 weeks post injection. (d) Whole brain EGFP fluorescence images (green) after intravenous injection of AAV9, PHP.eB, and CNSRCV300 at different doses, together with representative EGFP fluorescence images of the hippocampus, cortex, thalamus, striatum, medulla, and cerebellum. Scale bars are indicated in the first image of each brain region. (e) Representative EGFP fluorescence images of liver tissue after intravenous injection of AAV9, PHP.eB, and CNSRCV300 at different doses. Scale bars are shown in the first panel. (f) Quantification of EGFP positive cells after intravenous injection of AAV9, PHP.eB, and CNSRCV300 at different doses. Cortical EGFP positive cells were counted as representative of whole brain expression; for liver, EGFP positive cells were counted within a fixed size region (≈1.035 mm × 1.035 mm, corresponding to 1500 × 1500 pixels). Each dot represents one mouse; n = 3–4 per group. Statistical analysis was performed using two way ANOVA; *p ≤0.05, **p ≤ 0.01, ***p ≤ 0.001, ****p ≤ 0.0001.

PHP.eB is an adeno-associated virus (AAV) capsid variant selected via directed evolution using the M-CREATE methodology [11-13]. It demonstrates high efficiency in crossing the blood-brain barrier (BBB) and transducing murine brain cells, standing as one of the most comprehensively characterized variants derived from the AAV9 serotype. Studies have confirmed that PHP.eB-mediated BBB crossing is dependent on the vascular endothelial-specific receptor LY6A, which is abundantly expressed in C57BL/6J mice[6, 14]. However, the activity of LY6A is species-restricted; it is non-functional in other mouse strains, such as BALB/c, and lacks detectable activity in non-human primates (NHPs). This receptor specificity significantly limits the broader application of PHP.eB beyond C57BL/6J model[15, 16].

The capsid variant BI⍰hTFR1 was selected by targeting the human blood⍰brain barrier receptor TFRC and demonstrates a 40⍰ to 50⍰fold higher level of reporter gene expression in the central nervous system compared to AAV9 [10]. TFRC is abundantly expressed on the BBB surface in both humans and non-human primates, and its gene sequence is highly conserved. These features suggest that BI-hTFR1 may possess cross-species tropism and holds significant potential for advancing AAV capsid research toward clinical translation [10].

CNSRCV300 (also designated STAC-BBB; international patent WO2024238684A1) is a high-efficiency capsid variant developed by Sangamo Therapeutics based on directed evolution of AAV9, although the specific receptor responsible for its BBB crossing remains unidentified [17]. This variant has been reported to exhibit cross-species transduction capability, effectively targeting the brain across the BBB in C57BL/6J mice, NHPs, and human models. According to official data from Sangamo Therapeutics, CNSRCV300 demonstrates more than 65-fold higher transduction efficiency compared to AAV9 [18].

This study employed a classical adeno-associated virus (AAV) in vitro packaging system. The transfer plasmid, carrying the EGFP reporter gene, was driven by either the constitutive CAG promoter or the neuron-specific hSyn promoter. The helper plasmid pHelper and the packaging plasmid pRep/Cap provided the necessary genetic elements in trans for viral replication and capsid protein assembly. The successfully packaged AAV particles were administered to mice via tail vein injection. This approach aimed to evaluate the expression profiles of the aforementioned three AAV9-derived capsid variants across different mouse tissues, as well as their capability to cross the blood-brain barrier and transduce brain tissue, under the regulation of the distinct promoters.

## Result

### Expression of CAG-Promoter-Driven AAV-PHP.eB and AAV-CNSRCV300 in the Brains of C57BL/6J Mice

Both PHP.eB and CNSRCV300 are AAV capsids reported in the literature to mediate efficient transduction in the central nervous system of C57BL/6 mice. To systematically compare their tissue expression profiles *in vivo*, this study constructed the transfer plasmid pAAV⍰ CAG⍰ EGFP, in which EGFP expression is driven by the CAG promoter. The plasmid was packaged into the two aforementioned capsids. The resulting AAV particles were administered via retro ⍰ orbital sinus injection at two dose levels: 1.0×10^11^ vg per mouse and 4.0×10^11^ vg per mouse, with AAV9 included as a control (Fig. 1b, c). All mice remained in good general condition following injection. Four weeks later, brain, liver, and other tissues were collected for sectioning and subsequent analysis (Fig. 1c).

For both the PHP.eB and CNSRCV300 capsids, the EGFP they mediated exhibited a broader distribution pattern in the mouse brain compared to AAV9 (Fig. 1d). Relative to PHP.eB, CNSRCV300 showed lower infection efficiency in the cortex but demonstrated high-efficiency transduction in regions such as the hippocampal CA1 and DG areas, whereas the opposite trend was observed in the cerebellum (Fig. 1d). In this study, although the infection efficiency of CNSRCV300 did not reach the levels reported in the literature[18], the number of EGFP-positive cells in the cortical region was still approximately 7 times that of AAV9 (at a dose of 4.0×10^11^ vg: AAV9 mean = 193, CNSRCV300 mean = 1408). In the PHP.eB group, this value exceeded 40 times that of AAV9 (at the same dose: AAV9 mean = 193, PHP.eB mean = 8334) (Fig. 1f). In contrast to the results in brain tissue, the hepatic accumulation of CNSRCV300 was the lowest among the three capsids, being only one⍰fifth that of AAV9 or half that of PHP.eB (at a dose of 4.0×10^11^ vg) (Fig. 1e, f). This distribution pattern suggests that, compared to PHP.eB, CNSRCV300 may possess lower liver tropism and potentially reduced hepatotoxicity.

### Peripheral Organ Tropism of AAV-PHP.eB and AAV-CNSRCV300 Capsids

To investigate potential adverse effects associated with excessive peripheral expression, we collected selected peripheral organs and examined EGFP expression in the lung, heart, kidney, and spleen via tissue sectioning (Fig. 2). The results showed that upon high-dose administration, the CNSRCV300 capsid led to detectable expression in cardiac and lung tissues, while its expression was barely detectable in the kidney or spleen. In contrast, PHP.eB exhibited detectable expression signals in all four organs examined. This indicated that PHP.eB possessed a broader peripheral transduction tropism *in vivo* compared to CNSRCV300. Consequently, in studies involving high-dose AAV administration, CNSRCV300 presented a potentially more favorable safety profile than PHP.eB.

**Fig 2.**
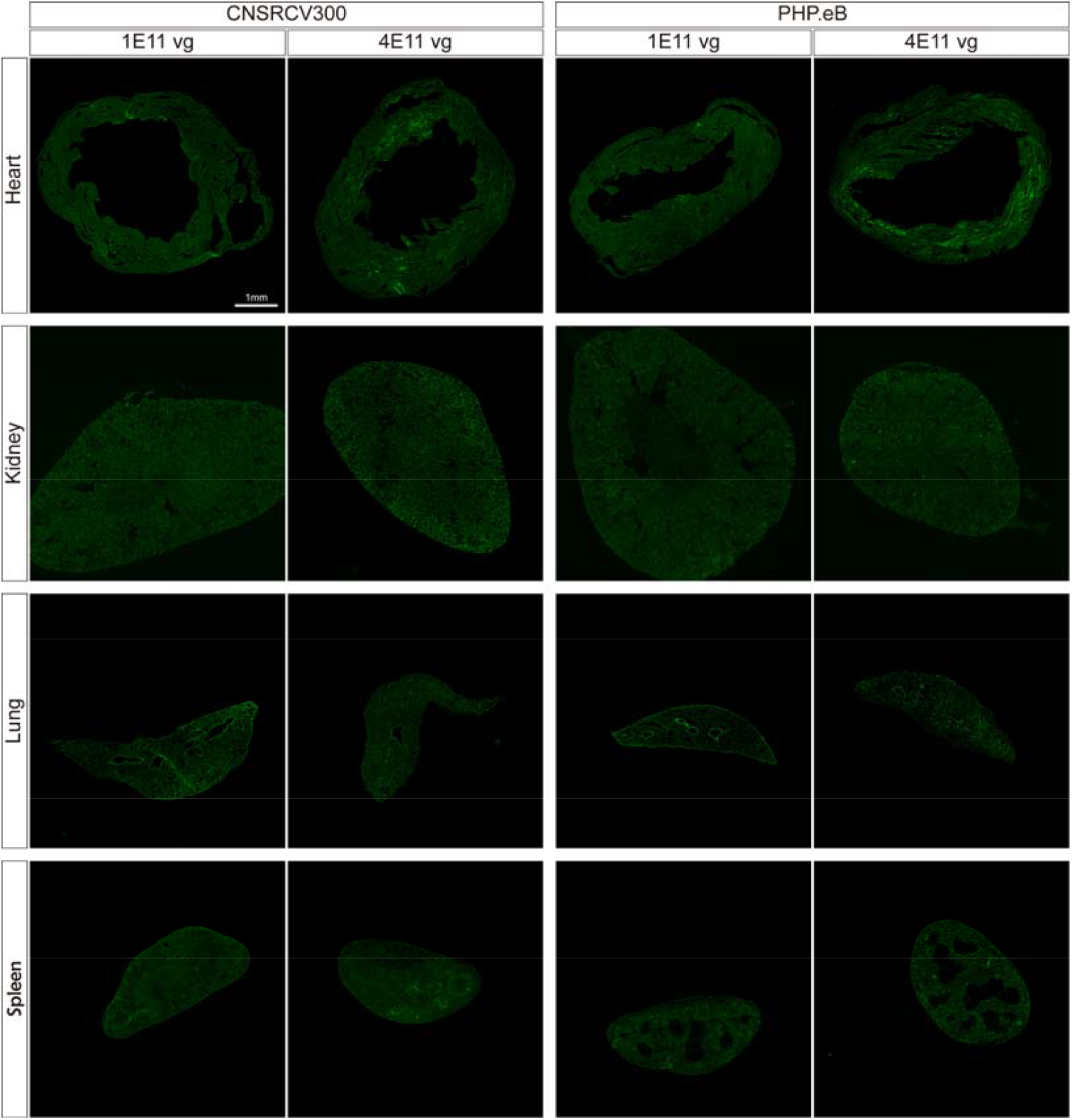
EGFP fluorescence expression in peripheral tissues of adult mice following intravenous injection of AAV-PHP.eB and AAV-CNSRCV300 carrying the CAG promoter. Adult C57BL/6J mice were intravenously injected with either AAV-PHP.eB or AAV-CNSRCV300 at doses of 1.0×10^11^ vg and 4.0×10^11^ vg. EGFP expression in peripheral tissues was examined 28 days post-injection. Representative fluorescence images of the lung, heart, kidney, and spleen are shown (n=4 per group). Scale bars are indicated in the images.

### AAV expression driven by the hSyn promoter exhibits neuron-specificity

The use of neuron-specific promoters is expected to restrict the off-target expression of AAV in peripheral organs. Therefore, we constructed the transfer plasmid pAAV-hSyn-EGFP, driven by the hSyn promoter, to achieve stable EGFP expression in neurons (Fig 3a, b). Additionally, this study included the AAV capsid variant BI-hTFR1—selected based on the human transferrin receptor (TFRC)—as a control.

**Figure 3.**
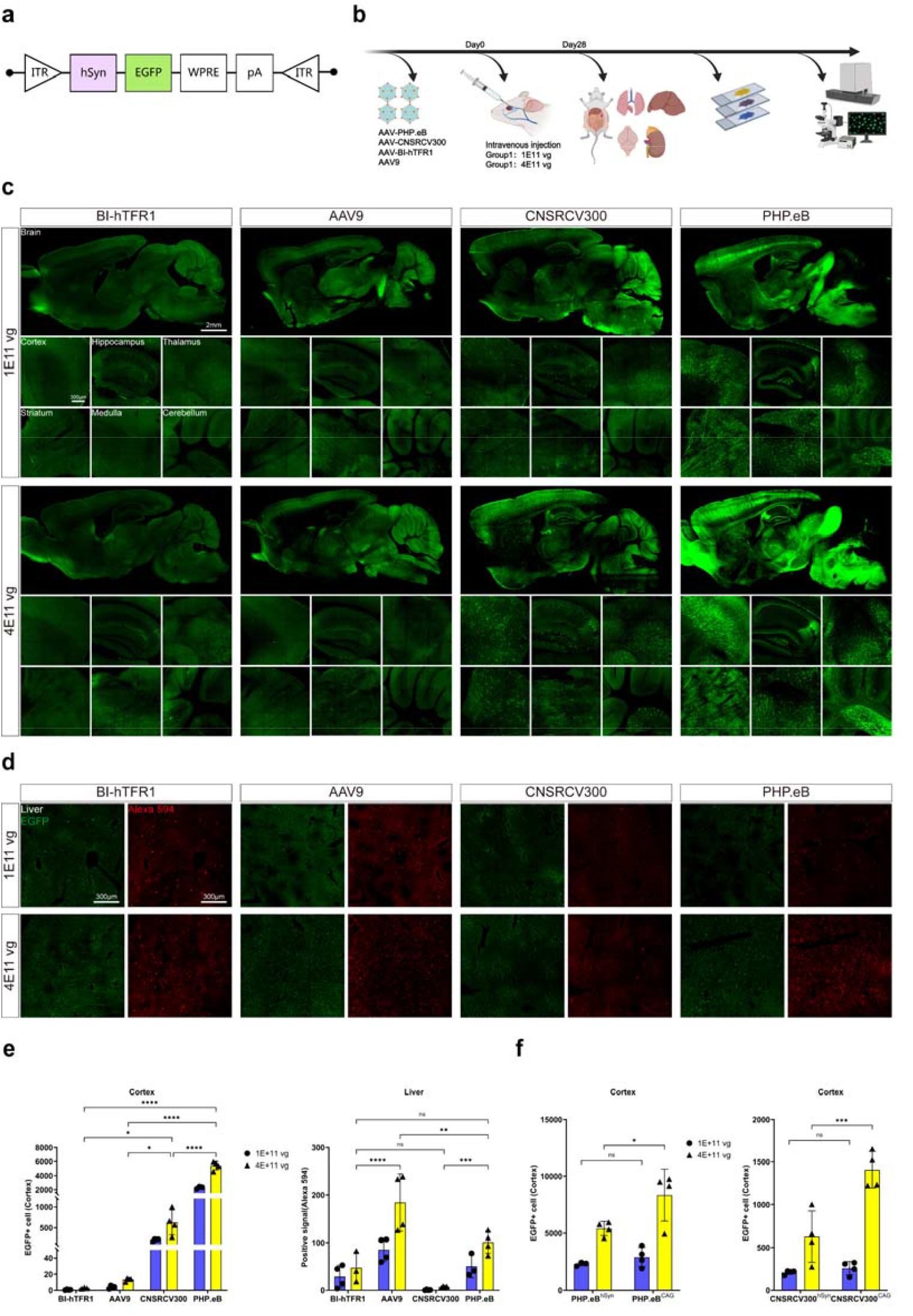
Transgene expression in brain and liver tissues mediated by AAV9-related capsids carrying the hSyn promoter. (a) Schematic diagram of the transgene plasmid structure containing the neuron-specific promoter hSyn. (b) Experimental procedure: Packaged AAV9, BI hTFR1, PHP.eB, and CNSRCV300 viruses (c) were intravenously injected into adult C57BL/6J mice at doses of 1.0×10^11^ vg and 4.0×10^11^ vg (n = 3–4 per group). Tissues were collected and sectioned for analysis 4 weeks post-injection. Panoramic views of whole-brain EGFP fluorescence expression (green signal for EGFP) after intravenous injection of BI hTFR1, AAV9, PHP.eB, and CNSRCV300 at different doses, along with representative fluorescence images of the hippocampus, cortex, thalamus, striatum, medulla, and cerebellum. Scale bars are indicated in the first image of each corresponding brain region. (d)Immunohistochemical detection of EGFP expression in the liver after intravenous injection of BI hTFR1, AAV9, PHP.eB, and CNSRCV300 at different doses. Green indicates EGFP fluorescence; red represents Alexa Fluor 594 immunostaining signal with anti-EGFP antibody. Scale bars are shown in the first panel. (e)Quantification of EGFP positive cells after intravenous injection of BI hTFR1, AAV9, PHP.eB, and CNSRCV300 at different doses. For brain tissue, EGFP positive cells in the cortical region (green signal) were counted as representative of whole-brain levels. For liver tissue, EGFP positive cells (red signal) were counted within a fixed-size area (approximately 1.6 mm × 1.6 mm) in immunostained images. Each dot represents an individual mouse, with n = 3–4 per group. (f)Comparison of EGFP expression in the mouse brain after infection with CNSRCV300 and PHP.eB under the control of either the CAG or hSyn promoter (using the cortical region as representative of whole brain, green signal). Each dot represents an individual mouse, with n = 3–4 per group. Statistical analysis was performed using two-way ANOVA; *p ≤ 0.05, **p ≤ 0.01, ***p ≤ 0.001, ****p ≤ 0.0001.

**Fig 4.**
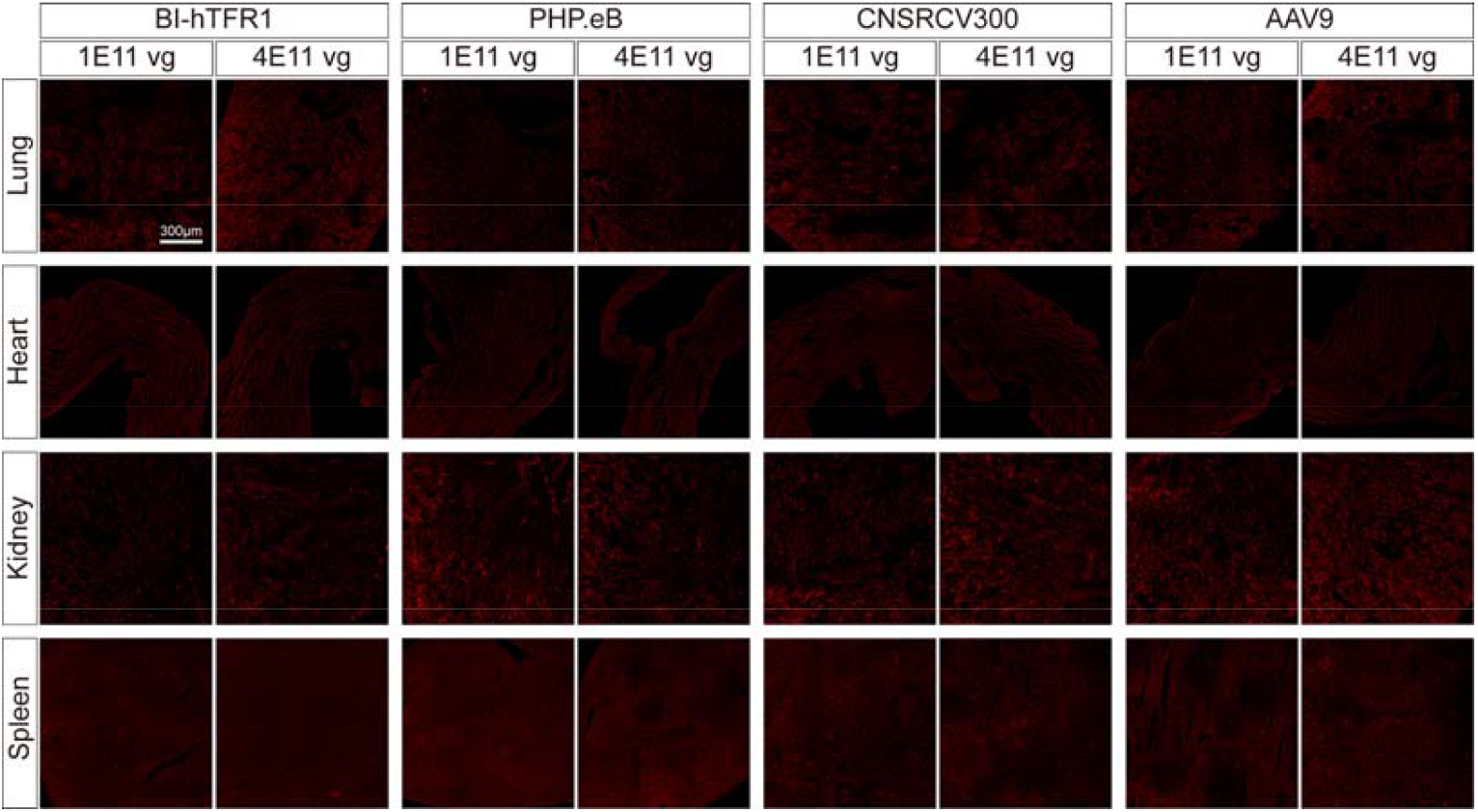
Immunohistochemical detection of AAV transduction in peripheral organs of adult mice using BBB ⍰ crossing neurotropic capsids. Mice were intravenously injected with the indicated capsids at doses of 1.0×10^11^ ⍰ vg or 4.0×10^11^ ⍰ vg. Twenty ⍰ eight days post ⍰ injection, EGFP expression was detected on tissue sections by immunohistochemistry (red signal). Representative images of the lung, heart, kidney, and spleen are shown (n = 3–4 per group). Scale bars are indicated in the images.

Fluorescence imaging of mouse brain tissue sections showed that the EGFP gene carried by all viral vectors was normally expressed in neurons, while its expression in other brain cell types, such as astrocytes, was significantly reduced (Fig. 3c). Based on the statistical data of cortical EGFP-positive cells, at high doses, PHP.eB exhibited levels more than 8 times that of CNSRCV300 (PHP.eB mean 5407, CNSRCV300 mean 626.6) (Fig 3e). Compared to the CAG group, reporter gene expression of PHP.eB and CNSRCV300 in astrocytes was hardly detectable in the hSyn group, and PHP.eB also exhibited downregulated reporter gene expression in cerebellar Purkinje cells. Moreover, CNSRCV300 maintained high expression in the hippocampal CA1 and DG regions, which was consistent with the observations in the CAG group (Fig. 1d and 3c). AAV-BI ⍰ hTFR1 exhibited low transduction efficiency in the mouse brain, a finding consistent with previous literature reports [10] (Fig. 3c).

Compared to the CAG promoter group, AAV expression in the liver was markedly decreased in the hSyn promoter group. At an injection dose of 1.0×10^11^ vg, the reporter gene expression levels of the AAVs were barely detectable, whereas at a dose of 4.0×10^11^ vg, apparent expression of the reporter gene from AAV9 and PHP.eB could be directly observed (Fig. 3d). Quantitative immunohistochemical analysis showed that these AAV particles were still present in the cells (Fig. 3d). In terms of liver accumulation, BI-hTFR1 showed a level similar to that of PHP.eB. Among the four AAV variants, CNSRCV300 still exhibited the lowest level of hepatic enrichment. At a high dose (4.0×10^11^ vg), its enrichment was only approximately 1/15 that of AAV9 (under identical measurement conditions: CNSRCV300 mean 7.09, AAV9 mean 184.2) (Fig. 3e). This trend aligns with the results observed in the previous CAG promoter group.

Studies have shown that single-amino-acid mutations in the AAV9 capsid can modulate its hepatotropism. For instance, the substitution of asparagine with aspartate at position 270 of the VP1 subunit (N270D) has been demonstrated to significantly reduce its accumulation in the liver [19]. Based on this, we introduced the N270D mutation into each of the four capsid variants used in this study and packaged them with the transfer plasmid pAAV-hSyn-EGFP to generate recombinant viruses (Fig S1a). This aimed to evaluate whether these engineered variants could maintain their original central nervous system transduction capability while reducing hepatotropism. Observation of viral expression via whole-brain tissue section scanning and quantitative analysis revealed that, regardless of the mutation’s effect on liver accumulation, all N270D-carrying AAV variants exhibited a marked decrease in both infection efficiency and transgene expression levels in the mouse brain (Fig. S1b).

In addition, we compared the brain expression profiles of two highly efficient capsids— AAV-CNSRCV300 and AAV-PHP.eB—under the control of either the CAG promoter or the hSyn promoter. Quantitative analysis showed that at a low dose (1.0×10^11^ ⍰vg), there was no significant difference in the number of EGFP-positive cells in the cortical region between the two capsids. However, at a high dose (4.0×10^11^ ⍰ vg), the CAG promoter group exhibited approximately twice as many positive cells as the hSyn promoter group (Fig.⍰3f). We speculate that at low doses, the overall transduction efficiency of AAV is limited, masking differences in expression driven by the two promoters, whereas at high doses, viral transduction approaches saturation, allowing the transcriptional characteristics of each promoter to become fully apparent. Fluorescence imaging further revealed a greater number of EGFP-positive astrocytes in the cortex of the CAG promoter group (Figs.⍰1d and ⍰3c). This observation supports the role of the promoter in shaping AAV tropism: unlike the broadly active CAG promoter, the neuron-specific hSyn promoter is inefficient at initiating transcription in non-neuronal cell types such as astrocytes and Purkinje cells, thereby conferring stricter cellular specificity.

The neuron-specificity of the hSyn promoter was also demonstrated in other tissues. In peripheral organs, even under a high dose (4.0×10^11^ ⍰ vg), EGFP expression was barely detectable by direct fluorescence microscopy. This does not, however, indicate a failure of AAV infection. Immunofluorescence staining revealed low levels of AAV-mediated expression in some peripheral organs (Fig.⍰4). These findings suggest that AAV particles may still reside within tissue cells, which could constitute a potential risk in clinical safety evaluation.

## Discussion

It has been reported that AAV9, PHP.eB, CNSRCV300, and AAV-BI-hTFR1 are all AAV capsid variants capable of crossing the blood–brain barrier and infecting the central nervous system [10, 11, 18]. In intravenous injection experiments performed in C57BL/6 mice, both AAV-PHP.eB and AAV-CNSRCV300 exhibited highly efficient BBB penetration and neuronal labeling, with cortical neuron transduction efficiencies increased by tens of times compared to AAV9. In contrast, BI-hTFR1 showed almost no effective neuronal infection (Fig. 3c), which is likely because BI-hTFR1 is an engineered capsid developed based on human TFRC and faces significant species-specific limitations, resulting in inefficient cross-species infection in mice.

The differences in the expression distribution of these AAV variants in the brain are generally attributed to their varying capacities to cross the blood-brain barrier (BBB), which is closely associated with their specific BBB receptors. AAV enters the brain via receptor-mediated transcytosis facilitated by specific receptors on the BBB. Several vascular surface receptors involved in this process have been identified, including LY6A[6, 15, 16, 20], LY6C1[15], CA-IV[14, 15], ALPL[21], TFRC[2, 8-10], among others. Most of these membrane proteins exhibit species specificity, meaning that AAV variants selected based on these receptors often face challenges in cross-species infection, thereby hindering their translation into clinical research. ALPL (also known as tissue-nonspecific alkaline phosphatase, TNAP) is a conserved membrane protein at the genetic level, sharing over 92% amino acid identity among humans, mice, and macaques [21]. The AAV9 variant VCAP-102, developed by Tyler C Moyer et al. [21], relies on ALPL to cross the BBB and has demonstrated strong affinity for ALPL in mice, pigs, humans, and macaques. This highlights the significant potential of ALPL in the development of engineered AAV capsids through receptor-based screening.

Specificity promoters do not affect the infection efficiency of AAV but can significantly reduce transgene expression in non⍰target cells. In this study, compared to the hSyn promoter, the CAG promoter resulted in more EGFP⍰positive signals in non⍰neuronal cell types such as glial cells and Purkinje cells. In peripheral tissues, AAV expression levels are jointly regulated by both the capsid type and the promoter. Taking PHP.eB as an example, AAV carrying the hSyn promoter showed markedly lower expression in peripheral tissues compared to the CAG group, to the extent that it was barely detectable directly; however, immunofluorescence staining still revealed a large number of infected cells (Fig. 3d). These infected cells may still pose potential safety risks to the organism. In contrast, although the brain-targeting efficiency of CNSRCV300 is slightly lower, its peripheral tropism is also significantly reduced, suggesting that it may offer better safety than PHP.eB in pharmacotoxicological evaluations. We attempted to engineer these capsids to further reduce hepatotropism, but this modification severely compromised their brain-targeting ability—a phenomenon that has not been systematically reported in studies [19].

Our study is limited to investigating the expression profiles of AAV capsids in mouse tissues and does not extend to elucidating the specific mechanisms underlying their traversal of the blood–brain barrier or to quantifying their potential peripheral toxicity. These aspects should be considered and addressed in future preclinical testing of novel AAV variants.

## Methods

### Animals

Adult male C57BL/6J mice (aged 8-10 weeks) were purchased from Vital River Laboratories (Beijing) Co., Ltd. The animals were housed under a 12-hour light/12-hour dark cycle with free access to standard diet and water. All animal experimental protocols were approved by the Ethics Committee of Lingang Laboratory. Prior to drug injection experiments, mice were acclimatized in group housing for at least one week.

### Plasmid

Based on the parental plasmid pUCmini⍰iCAP⍰PHP.eB (Addgene No. 103005), we constructed the following three capsid expression plasmids: first, the peptide sequence DGTLAVPFK after residue 586S of the PHP.eB capsid was replaced with AQYSRIGPN to obtain the BI⍰hTFR1 capsid expression plasmid; second, the same peptide sequence was replaced with AQ to restore the original AAV9 capsid sequence; finally, the peptide sequence YVNIMDDMD was inserted at residues 588Q and 589A of the AAV9 capsid to generate the capsid variant CNSRCV300. All constructed capsids carry a common mutation relative to wild⍰type AAV9 (K449R, based on the NNK library), which has been reported to have no significant impact on viral packaging or transduction efficiency[17, 18]. In the specific experimental procedure, we first performed double digestion of the plasmid pUCmini-iCAP-PHP.eB using the restriction enzymes BsiWI and AgeI. Subsequently, using this plasmid as a template, PCR amplification was carried out with a high-fidelity DNA polymerase. All amplification reactions used the same forward primer (5⍰-GACTCAGACTATCAGCTCCCGTACGTGCTCGGGTCGG-3 ⍰), while reverse primers were individually designed according to the different target fragments (sequences are listed below). Finally, the digested products and PCR products were assembled seamlessly using NEBuilder^®^ HiFi DNA Assembly Master Mix to obtain the recombinant plasmid.

Reverse primers were used for:

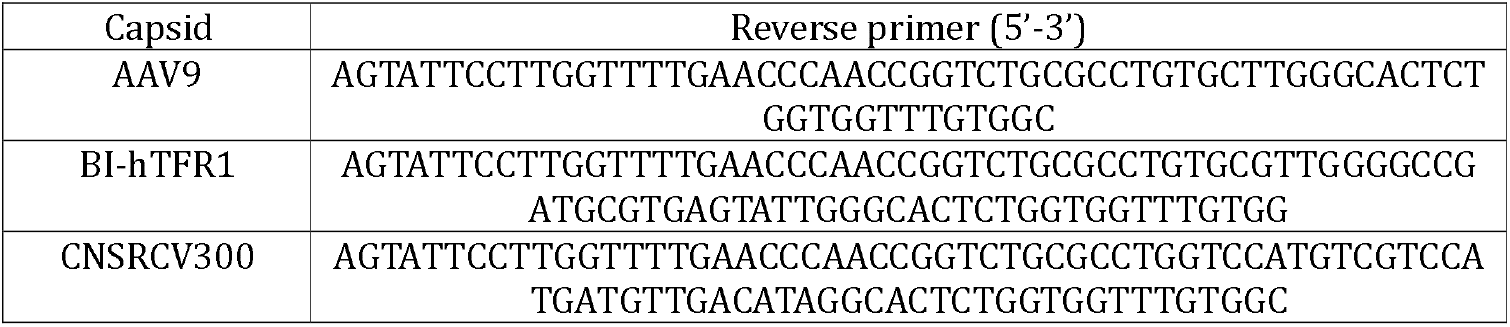

### AAV Production

HEK293T cells were seeded in 150⍰mm dishes coated with poly-L-lysine and cultured at 37 ⍰ ⍰ under 5⍰% CO_2_ until they reached 80–90 ⍰ % confluence. Transfection was performed using polyethylenimine (PEI) with a mixture of the helper plasmid pHelper, the packaging plasmid pRep/Cap, and the transfer plasmid carrying the EGFP reporter gene. Culture supernatants and cell pellets were collected separately at 72 ⍰ h and 120 ⍰ h post-transfection. Viral particles in the supernatant were precipitated by adding 40 ⍰ % polyethylene glycol ⍰ 8000 (PEG ⍰ 8000) at a 1:5 (v/v) ratio. To release intracellular virus, the cell pellets were subjected to repeated freeze-thaw cycles combined with Benzonase treatment, followed by centrifugation to remove cellular debris. The clarified supernatant was then purified by iodixanol density-gradient ultracentrifugation. The purified virus was buffer-exchanged into DPBS and concentrated using an ultrafiltration device.

Viral titers were determined by quantitative polymerase-chain reaction (qPCR) targeting the WPRE sequence within the viral genome, and these values served as the basis for subsequent animal injection schemes. Primers were used for:

WPRE forward primer (5’-3’): CCCGTATGGCTTTCATTTTCTCC

WPRE reverse primer (5’-3’): GGCAATGCCCCAACCAGTG

WPRE probe: FAM-TGGTGTGCACTGTGTTT-MGB

### Intravenous Injection

AAV was administered via retro-orbital venous plexus injection. Mice were anesthetized with isoflurane, and a diluted AAV solution (100⍰μL per mouse) was injected into the retro-orbital venous plexus using an insulin syringe. In this study, two dose groups were used: 1.0×10^11^ ⍰ vg and 4.0×10^11^ ⍰ vg per mouse, respectively.

### Animal Tissue Processing

Perfusion was performed four weeks after AAV injection. Following deep anesthesia, mice underwent transcardiac perfusion, initially with phosphate-buffered saline (PBS) followed by fixation with 4% paraformaldehyde (PFA). Brain tissue and peripheral organs were then harvested and post-fixed overnight in 4% PFA at 4°C. After fixation, brain tissue was sectioned directly along the sagittal plane at 50 ⍰ μm thickness using a vibrating microtome (Leica). Peripheral organs were cryoprotected overnight in 30% sucrose PBS solution at 4°C until the tissues sank, then embedded in OCT compound and sectioned at 16 ⍰ μm thickness using a cryostat (Leica).

### Imaging Analysis

Tissue sections from the brain and peripheral organs were scanned using a whole-slide imaging system (VS200, Evident) to obtain EGFP fluorescence images. The number of EGFP-positive cells in cortical regions and liver tissues was quantified using ImageJ software.

### Immunohistochemistry

Tissue sections were permeabilized with PBS containing 1% Triton X-100 for 15 min at room temperature, followed by blocking with 3% bovine serum albumin (BSA) for 1 h at room temperature. The sections were then incubated overnight at 4 ⍰ °C with the primary antibody (chicken anti-GFP, Aves Labs, GFP-1020; diluted 1:500). After three washes with PBS, the sections were incubated for 1 h at room temperature in the dark with the secondary antibody (donkey anti-chicken Alexa Fluor 594, Jackson ImmunoResearch; diluted 1:400). Finally, nuclei were counterstained with DAPI.

The stained sections were imaged using a confocal microscope (Stellaris ⍰ 5, Leica). Images were acquired and initially processed with the Las ⍰ X imaging software (Leica). Alexa Fluor ⍰ 594-positive cells in liver sections were quantitatively counted using ImageJ software.

### Statistical Analyses

All quantitative data are expressed as the mean ± standard deviation (mean ± SD). Statistical analysis was performed using GraphPad Prism software (version 9.5.0). Differences between groups were assessed by two-way analysis of variance (two-way ANOVA). A P value of < 0.05 was considered statistically significant.

## Acknowledgments

This study was supported by the Lingang Laboratory Project (LG-GG-202401-01-ADA070400). We thank Xueping Gao and Xiao Wang for their assistance in the AAV production process. We also thank Xianda Wang for his help with data processing and image processing methods.

## Author Contributions Statement

Conceptualization: JZ, LZ, LJ; Experimentation: JZ, XG, MS, WL; Resources: XZ; Data analysis: JZ; Manuscript preparation: JZ, LZ, LJ.

## Data Transparency Statement

Data, analytic methods, and study materials will be made available to academic investigators upon reasonable request.

