## Supplemental Figure 1 for "Comparative Study of BBB-Targeting AAV Capsids on Central Nervous System Delivery Efficiency" (from another version of this preprint: version 1, 2025-12-25)

**The PDF file includes:**

Supplemental Methods, Supplemental figures.

**Supplemental Methods**

**Single Amino Acid-Mutated Capsid Plasmid Construction**

In this study, based on the previously used capsid plasmids AAV9, BI‑hTFR1, CNSRCV300, and PHP.eB, we introduced a point mutation (N→D) at amino acid position 270 of the capsid protein coding sequence. The specific construction method was as follows: first, the original plasmids were double‑digested with the restriction enzymes AflII and BsiWI; then, a synthesized DNA duplex fragment containing the target mutation site (changing codon AAT for asparagine to GAC for aspartic acid at position 270) (synthesized by Genewiz) was assembled with the digested vector using the NEBuilder® HiFi DNA Assembly Master Mix system via homologous recombination. Finally, all four capsid plasmids carrying the N270D mutation were successfully obtained for subsequent experiments.

**Supplemental figures**

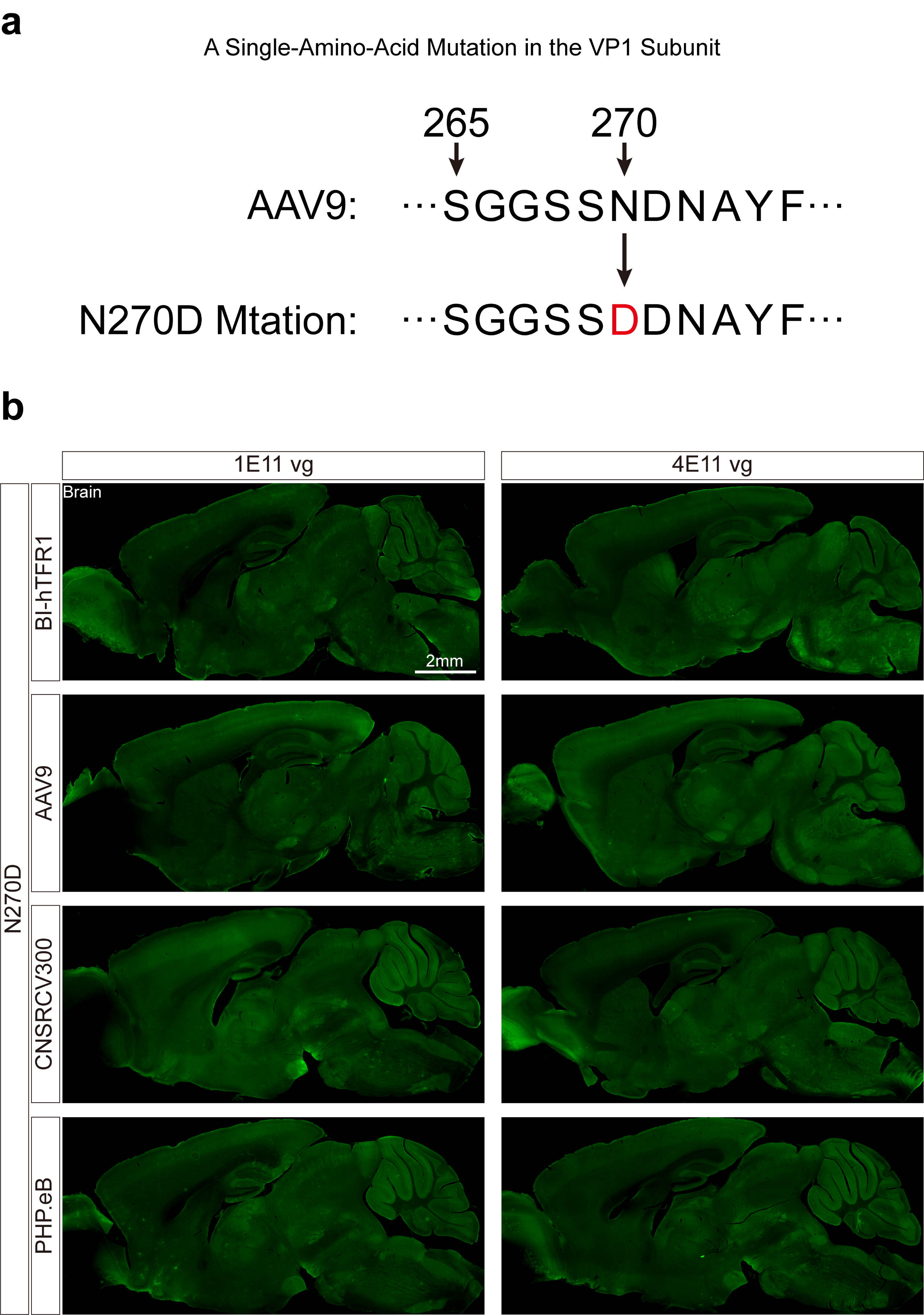

Fig S1. Brain transduction of AAV capsids following single‑amino‑acid mutation in the VP1 subunit of neurotropic variants

(a) Schematic of the single amino acid substitution introduced at residue 270 (N→D) in the VP1 subunit of an AAV9 derived neurotropic capsid.

(b) After packaging into viral particles, the mutant capsid was intravenously injected into adult C57BL/6J mice. EGFP reporter expression in the brain was assessed 28 days post injection. Representative images of EGFP fluorescence (green signal) in brain sections are shown (n = 4 per group). Scale bars are indicated in the images.
